# Characterization of the Root-Associated Microbiome Provides Insights into Endemism of *Thymus* Species Growing in the Kazdagi National Park

**DOI:** 10.1101/2023.03.30.534539

**Authors:** Gökçe Ercan, Muzaffer Arıkan, İ. Sırrı Yüzbaşıoğlu, F. Elif Çepni Yüzbaşıoğlu

**Affiliations:** Department of Molecular Biology and Genetics, Institute of Graduate Studies in Sciences, Istanbul University, Istanbul, Türkiye; Department of Medical Biology, School of Medicine, Istanbul Medipol University, Istanbul, Türkiye; Regenerative and Restorative Medicine Research Center (REMER), Research Institute for Health Sciences and Technologies (SABITA), Istanbul Medipol University, Istanbul, Türkiye; Department of Biology, Faculty of Science, Istanbul University, Istanbul, Türkiye; Department of Molecular Biology and Genetics, Faculty of Science, Istanbul University, Istanbul, Türkiye

**Author notes:** Corresponding author: F. Elif Çepni Yüzbaşıoğlu, Department of Molecular Biology and Genetics, Faculty of Science, Istanbul University, 34134 Vezneciler, Istanbul, Turkey. These authors contributed equally to this work (Gökçe Ercan, Muzaffer Arıkan).

**Keywords:** plant endemism, *Thymus*, rhizosphere, soil, microbiome, amplicon sequencing

## Abstract

Plant associated microbiomes have a large impact on the fitness of the plants in the particular environmental conditions. The root associated microbiomes are shaped by the interactions between the microbial community members, their plant host, and environmental factors. Hence, further understanding of the composition and functions of the plant root associated microbiomes can pave the way for the development of more effective conservation strategies for endangered endemic plants. Here, we characterized the bacterial and fungal microbiomes in bulk and rhizosphere soil of an endemic and a non-endemic *Thymus* species from Kazdagi National Park, Türkiye, *Thymus pulvinatus* and *Thymus longicaulis* subsp. *chaubardii*, respectively, by 16S rRNA gene and ITS amplicon sequencing. Our findings revealed no significant differences in alpha diversity between plant species and soil types. However, we found that the bacterial microbiome profiles differentiate not only *Thymus* species but also soil types while fungal microbiome profiles show distinct profiles particularly between the species in beta diversity. *Proteobacteria, Actinobacteria, Acidobacteria*, and *Chloroflexi* members form the core bacterial microbiome while the fungal core microbiome consists of *Ascomycota* and *Basidiomycota* members in both *Thymus* species. Moreover, we identified the association of the bacterial taxa contributing to the biogeochemical cycles of carbon and nitrogen and providing the stress resistance with the rhizosphere soil of endemic *T. pulvinatus*. In addition, functional predictions suggested distinct enriched functions in rhizosphere soil samples of the two plant species. Also, employing an exploratory integrative analysis approach, we determined the plant species-specific nature of transkingdom interactions in two *Thymus* species.

## 1. Introduction

Plants are key components of global biodiversity and essential for ecosystem sustainability; however, thousands of plant species face an increased risk of extinction worldwide (1). Endemic plants which thrive only in a specific geographic area are more prone to extinction (2) thus require particular attention when developing and establishing plant protection strategies. Clearly, a better understanding of the mechanisms of plant survival and growth is of paramount importance for successful biodiversity conservation.

Plants host diverse but taxonomically structured communities of microorganisms that colonize different plant tissues and the surroundings and play essential roles in plant health and growth (3). The rhizosphere is the zone of soil around the plant roots and composed of several soil microorganisms (4). Rhizosphere-associated microbiota play an important role in plant health, nutrient uptake, secondary metabolite production, immunity, and stress tolerance (5). Recent studies suggest a significant association between the plant microbiome and endemism. One of these studies focused on *Anthurium* species and found that the endemic species had special bacterial communities that supported the plant compared to other species (6). In another study, endemic *Hoffmannseggia doellii* growing at an altitude of 2800-3600 m in Atacama Desert was reported to have more diverse microbial communities compared to soils of other plants in the same region and bulk soil (7).

Approximately 11707 vascular plant species are known to grow in Türkiye, 3649 of which are endemic species (8). Kazdagi, a national park and one of the important biodiversity centers of Turkey, hosts about 800 plant species, 30 of which are endemic. There has been extensive research on the biodiversity and endemic plants in Kazdagi (9–11). However, to our knowledge, there is no study focusing on the potential relationship between plant microbiome and endemism for endemic plant species in Kazdagi.

In this study, we collected rhizosphere soil (RS) and bulk soil (BS) samples for two *Thymus* species, namely *T. longicaulis* subsp. *chaubardii* and *T. pulvinatus*. Although growing in the same location and environmental conditions with the other in Kazdagi, *T. pulvinatus* is a local endemic species. By applying 16S rRNA gene and internal transcribed spacer (ITS) sequencing to identify both bacterial and fungal microbiomes in soil samples from both species, we tested the following hypotheses: i) RS of endemic *T. pulvinatus* hosts a distinct bacterial and fungal community composition as a potential contributor to its endemism. ii) Functional profile differences of bacterial microbiome between two *Thymus* species are involved in different survival characteristics. iii) Transkingdom interactions differ between endemic and non-endemic *Thymus* species.

## 2. Materials and Methods

### 2.1. Sampling Area and Sample Collection

Sampling was carried out on August 29^th^, 2022, in Kazdagi National Park (Edremit, Balıkesir). Plant species identification of the materials was performed by Dr. Sirri Yüzbaşioğlu. The voucher specimens were preserved at the Herbarium of the Faculty of Pharmacy (ISTE), Istanbul University. *T. longicaulis* subsp. *chaubardii* samples (ISTE1183756) were collected near Kapidag watchtower stairs (39° 40’
s 53’’N - 26° 54’ 58’’E) at 1360 m height while *T. pulvinatus* samples (ISTE118375) were collected below Kapidag watchtower (39° 40’ 55’’N - 26° 54’ 55’’E) at 1350 m height. Biological triplicates (different plants) of each plant species were selected for the study. Rhizosphere soil (loosely attached to the plant roots) and bulk soil (surrounding the plant roots) were collected from 5-10 cm depth for each plant. Samples were stored at 4°C during the transfer to the laboratory, and frozen at -80°C when arrived at the laboratory.

### 2.2. DNA Extraction

DNA extractions were performed using QIAGEN DNeasy PowerSoil Pro Kit (Qiagen, Hilden, Germany) with modifications to the manufacturer’s protocol. In brief, to obtain rhizosphere DNA, plant roots with the attached soil were placed on sterile filter paper, cut into small pieces, and then suspended in 5 mL of filter-sterilized PBS. After 2 minutes of vortex, roots were removed and 1000 µL of the liquid sample was transferred to the PowerBead Pro Tube. Next, samples were centrifuged at 15.000 x g for 1 minute and supernatant discarded, which was repeated three times. After that, the manufacturer’s protocol was followed without any modification. DNA samples were stored at -20 °C until library preparation.

### 2.3. Library Preparation and Amplicon Sequencing

16S rRNA and ITS gene sequencing were performed in a two-step PCR amplification protocol. In 16S rRNA gene amplifications, universal bacterial primers F-5’-CCTACGGGNGGCWGCAG-3’ and R-5’GACTACHVGGGTATCTAATCC-3’ were used targeting V3-V4 region of 16S rRNA gene (12) while ITS1F (5’-CTTGGTCATTTAGAGGAAGTAA-3’) and ITS2R (5’-GCTGCGTTCTTCATCGATGC-3’) primers (13) were used for the investigation of fungal diversity. Amplicon libraries were prepared by following Illumina’s 16S rRNA metagenomic sequencing library preparation and fungal metagenomic sequencing protocol. MiSeq platform and 2×300 paired end sequencing kit were used for amplicon sequencing. A total of 24 amplicon libraries were sequenced, along with an extraction negative control and 2 no-template PCR controls.

### 2.4. Bioinformatics and statistical analyses

Paired end reads were demultiplexed based on their unique barcodes. Trimmomatic (14) was used for primer and barcode trimming and quality filtering steps. Filtered and quality checked paired end sequences were merged using FLASH (15). DADA2 pipeline (16) was used for taxonomic assignments and generation of ASV abundance tables. SILVA (v138) (17) and UNITE (v8.3) (18) databases were employed for bacterial and fungal communities, respectively. Potential contaminant sequences in the samples were filtered by the decontam (19). Only ASVs present in at least 2 samples and assigned at phylum level were included in the downstream analyses. Samples were rarefied to minimum sampling depth before performing alpha and beta diversity analyses which were performed using Phyloseq (20). Principal coordinate analysis (PCoA) using a dissimilarity matrix based on Euclidean distance was applied to examine the variation between samples in bacterial and fungal community compositions. Differentially abundant taxa between sample groups were examined using LefSe (21). Functional profiles of bacterial communities in samples were predicted using the Tax4Fun2 (22) and differentially abundant KEGG functional genes were determined using LEfSe. Data integration analysis using latent components (DIABLO) (23) was used to integrate 16S rRNA gene and ITS sequencing results and perform an exploratory examination of transkingdom interactions. Pheatmap (24) was used for construction of heatmaps using Euclidean distance as the similarity measure and clustering samples based on the “complete” method. The ggplot2 (25) was used for visualizations.

Statistical analyses were conducted in R 3.6.1. A Kruskal-Wallis test was used for alpha diversity comparisons. Permutational multivariate analysis of variance (PERMANOVA) from the vegan package was used for beta diversity comparisons. Differential abundance analyses of bacterial and fungal communities were performed with Linear discriminant analysis Effect Size (LEfSe) and tested using Kruskal–Wallis test and using Linear Discriminant Analysis (LDA) as implemented in LefSe.

### 2.5. Data availability

The raw 16S rRNA gene and ITS amplicon sequencing data produced in this study have been deposited in the NCBI Sequence Read Archive database, accession no. PRJNA943177.

## 3. Results

Soil samples were collected in the Edremit province of Türkiye (see “Methods” section). Both rhizosphere soil samples (named as LR and PR for *T. longicaulis* subsp. *chaubardii* and *T. pulvinatus*, respectively) and bulk soil samples (named as LS and PS for *T. longicaulis* subsp. *chaubardii* and *T. pulvinatus*, respectively) were obtained. Three samples were collected for each group which yielded a total number of 12 samples. Both 16S rRNA gene and ITS amplicon sequencing based microbiome analysis were performed for all soil samples (Figure 1). A total of 1.141.078 paired end bacterial amplicon sequences from the V3-V4 region of 16S rRNA gene and 1.568.574 paired end fungal amplicon sequences from the ITS region were obtained from all amplicon libraries. The 16S rRNA gene and ITS sequences generated 4664 and 3335 amplicon sequence variants (ASVs), respectively. The 16S rRNA gene ITS samples were rarefied to minimum sampling depth, 4085 and 15310 reads, respectively. After decontamination, filtering, and rarefaction, 1083 bacterial and 829 fungal ASVs were used for downstream analyses.

**Figure 1.**
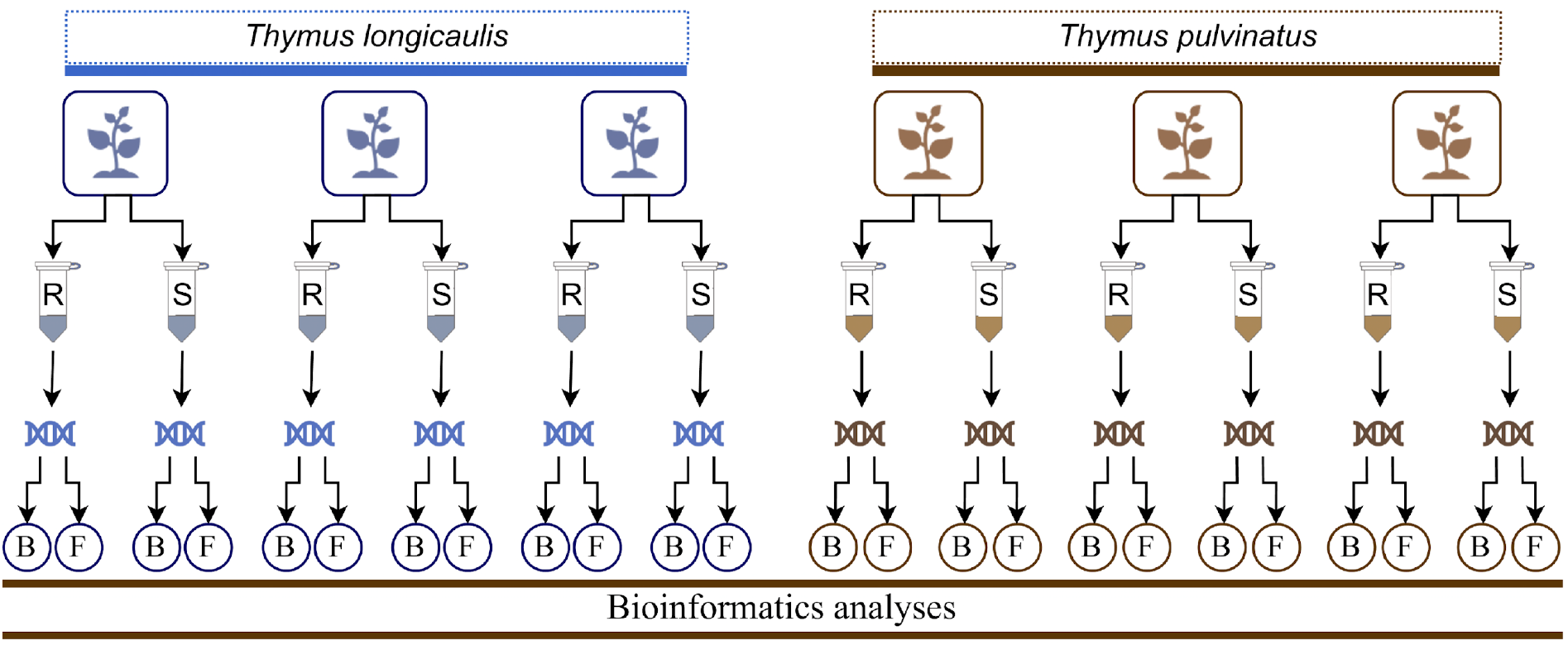
Experimental overview. Rhizosphere (R) and bulk soil samples (S) were collected from 3 plants for each *Thymus* species. Then, DNA extractions were performed for each soil sample. Next, bacterial (B) and fungal (F) microbiomes were profiled by amplicon sequencing. Finally, soil microbiome data was investigated using bioinformatics analyses.

### 3.1. Bacterial and fungal microbiome compositions

The bacterial ASVs assigned at phylum level showed that 10 most abundant phyla were *Proteobacteria, Actinobacteria, Acidobacteria, Chloroflexi, Bacteroidetes, Myxococcota, Verrumicrobia, Gemmatimonadetes, Planctomycetes* and *Firmicutes* (Figure 2A). Among these phyla, *Proteobacteria* (35%), *Actinobacteria* (33%), *Acidobacteria* (9%) and *Chloroflexi* (9%) were dominant across all samples on average. For fungi, 10 most abundant phyla included *Ascomycota, Basidiomycota, Mortierellomycota, Chytridiomycota, Mucoromycota, Olpidiomycota, Glomeromycota, Rozellomycota, Monoblepharomycota, Kickxellomycota* and *Aphelidiomycota*. Taxonomic analysis of the fungal community showed that on average, *Ascomycota* (72%) and *Basidiomycota* (19%) were dominant across all samples (Figure 2B).

**Figure 2.**
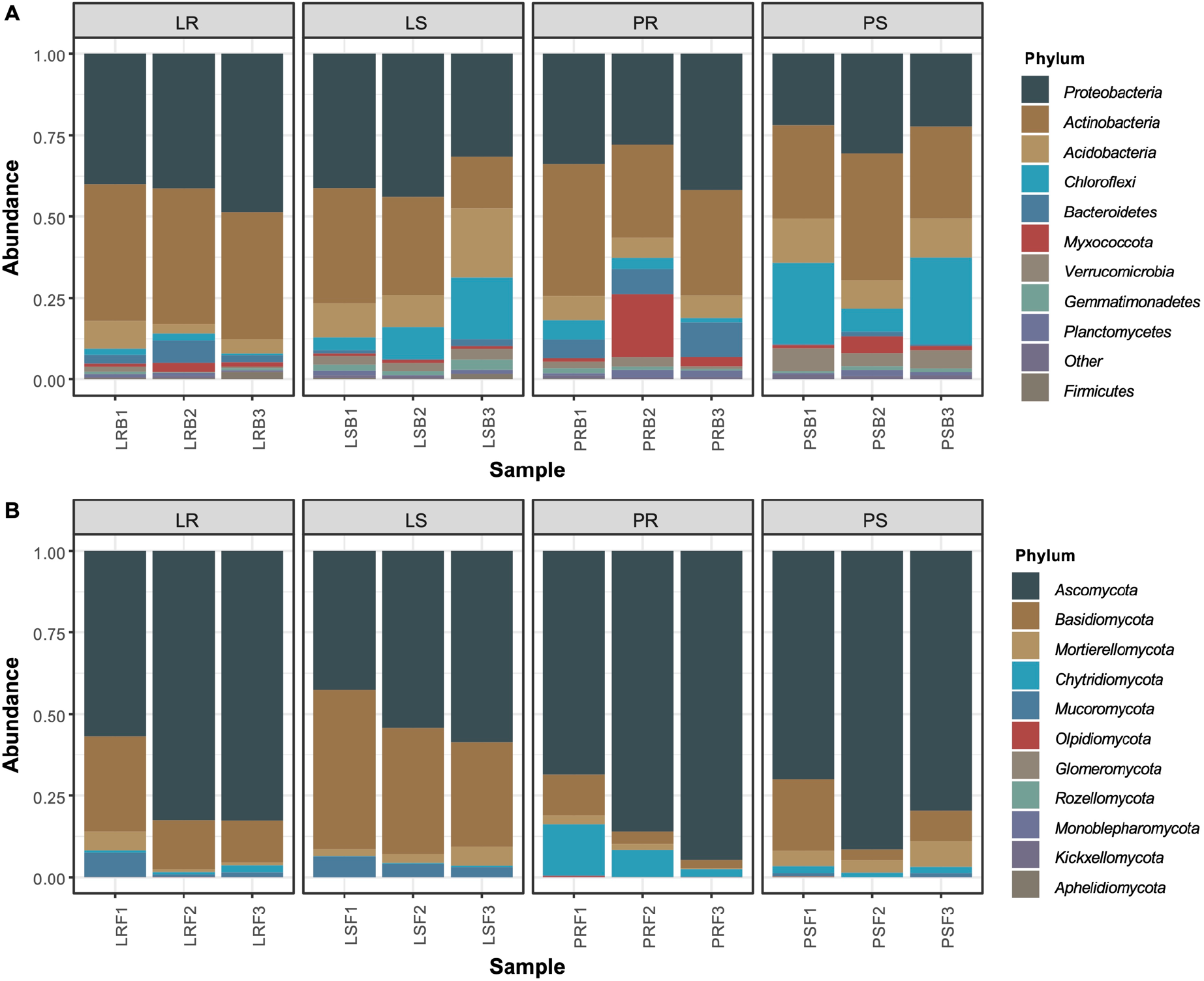
The top 10 most common bacterial (A) and fungal (B) phyla in soil microbiome samples. Phyla that were not among 10 most common taxa were grouped into “Other”. Each bar represents relative abundance distribution for a sample.

### 3.2. Structural diversity measures

Bacterial and fungal communities were evaluated using richness and diversity indices. There were no significant differences in alpha diversity indices (Chao1, Shannon, InvSimpson, Fisher) between *Thymus* species or rhizosphere and bulk soil samples for bacterial (Figure 3A) and fungal microbiomes (Figure 3B).

**Figure 3.**
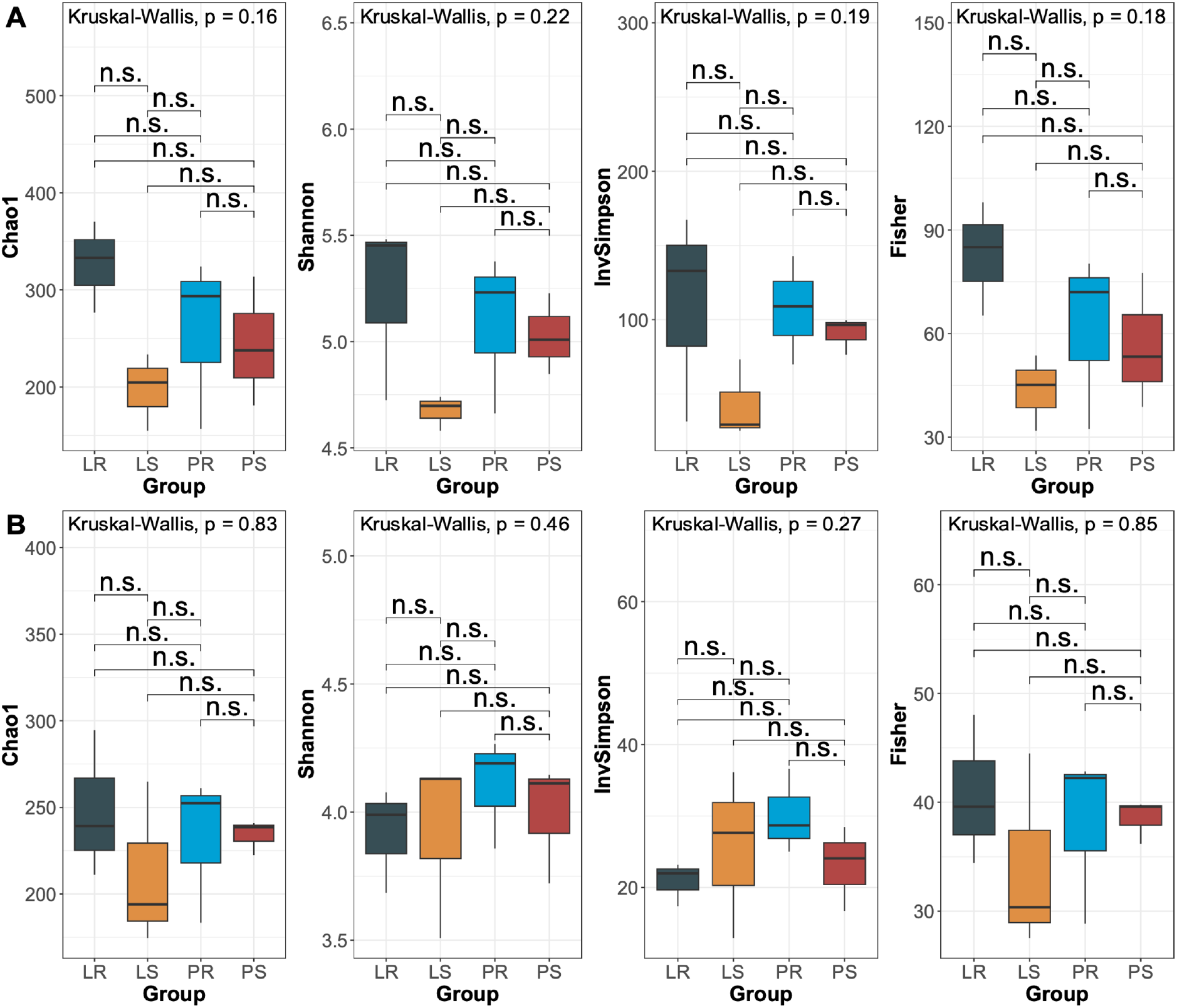
Alpha diversity (Chao1, Shannon, InvSimpson, Fisher) comparisons of bacterial (A) and fungal microbiome samples between study groups. Median estimates compared across study groups using the Kruskal-Wallis test. Boxes represent the interquartile range, lines indicate medians, and whiskers indicate the range. n.s: not significant.

To determine the variation between samples, PCoA using a dissimilarity matrix based on Euclidean distance was applied where axis 1 and axis 2 explained 52.1% and 50.6% variances among four sample types for bacterial and fungal microbiomes, respectively (Figure 4A and B). Bacterial microbiome samples clustered clearly according to not only the *Thymus* species but also rhizosphere and bulk soil types. Fungal microbiome samples clustered according to the *Thymus* species but separation between rhizosphere and bulk soil samples was not clear. PERMANOVA was used to test whether the samples cluster beyond what was expected by sampling variability. The results showed a significant difference between four sample types for both bacterial microbiome and fungal microbiome.

**Figure 4.**
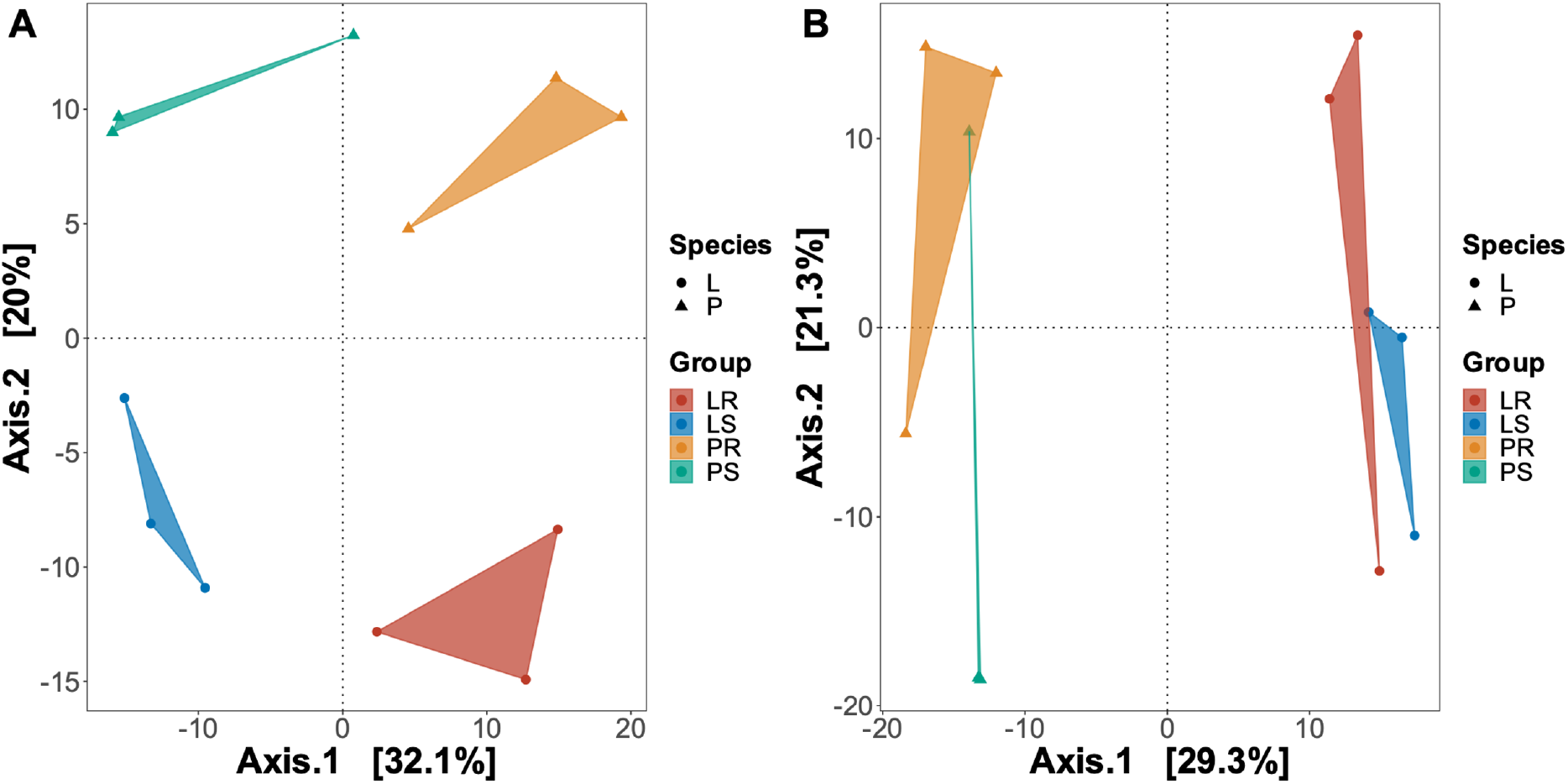
Beta diversity comparisons of bacterial (A) and fungal (B) microbiome samples between study groups. PCoA was calculated using euclidean distance. Color is indicative of the study group. Shape is indicative of *Thymus* species (L for *T. longicaulis* subsp. *chaubardii* and P for *T. pulvinatus*).

### 3.3. Differential abundance analysis

To determine which microbial taxa were significantly associated with sample groups, we performed differential abundance using LEfSe. For the bacterial community, we identified 20 differentially abundant bacterial genera with LDA greater than 4 enriched in different study groups (Figure 5A). Five genera were enriched in the rhizosphere soil of *T. longicaulis* subsp. *chaubardii*, including *Pseudomonas, Mycobacterium, Nocardioides, Streptomyces* and *Kribella* while rhizosphere soil of endemic *T. pulvinatus* included enriched unclassified *Acetobacteraceae, Sphingomonas, Blastocatella* and *Marmoricola*. For the fungal community, we identified 7 fungal genera with LDA greater than 4 enriched in LS and PR groups (Figure 5B). Among these, 5 genera (*Saitozyma*, unclassified *Tylosporaceae, Geminibasidium, Solicoccozyma*, unclassified *Leotiomycetes*) were enriched in bulk soil samples of *T. longicaulis* subsp. *chaubardii* (LS) while 2 genera (*Comoclathris* and unclassified *Chaetothyriales*) were enriched in the rhizosphere soil of endemic *T. pulvinatus*.

**Figure 5.**
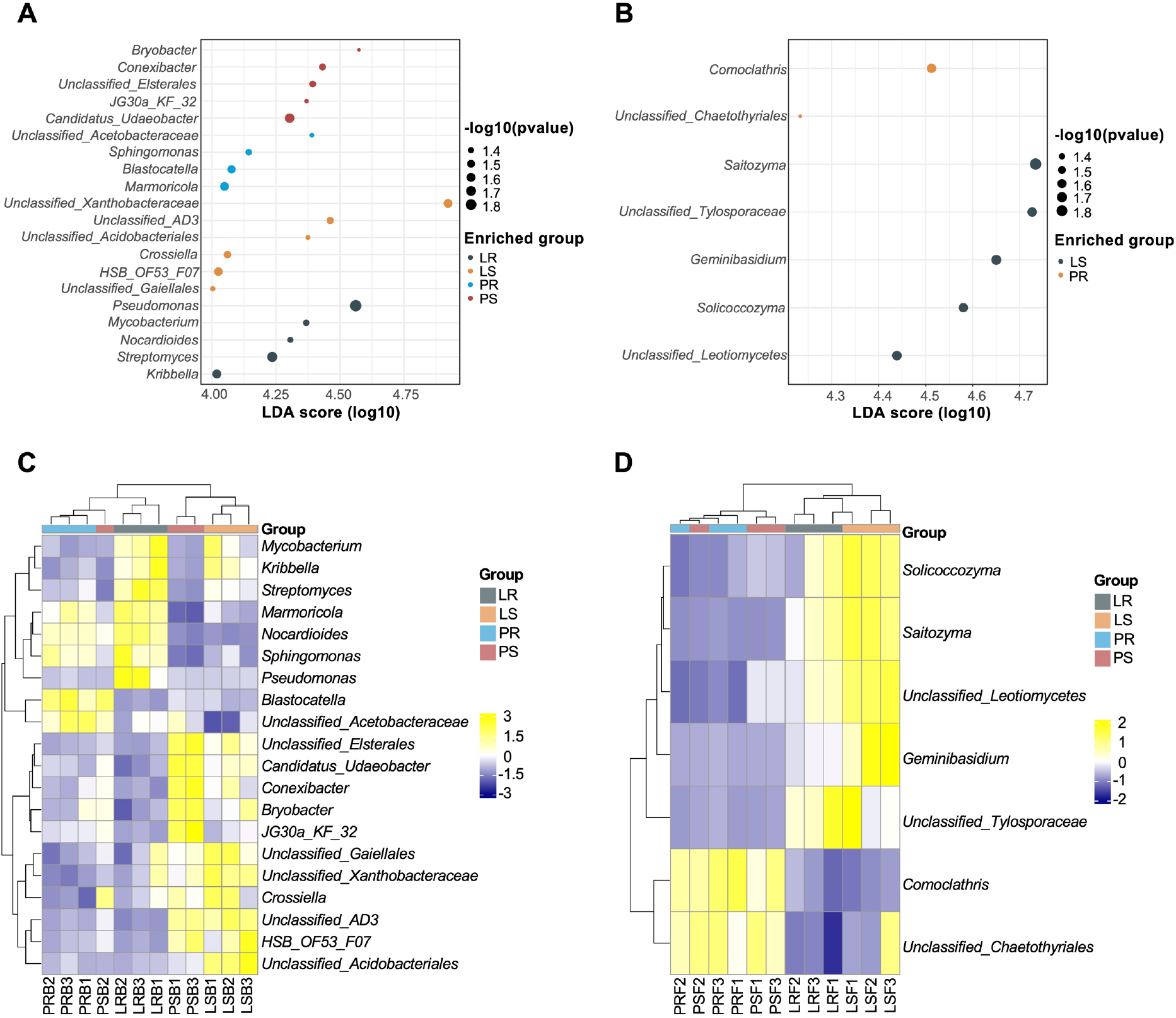
Abundance distribution of differentially abundant bacterial (A) and fungal (B) genera in the soil samples detected by LEfSe. LDA effect size (LEfSe) was calculated using LDA with p-value cutoff = 0.05 with LDA score > 4 of the genera. Dot sizes are indicative of -log10 (p value) while color is indicative of the study group with the enriched differentially abundant genera. Heatmap and hierarchical cluster analysis of bacterial (C) and fungal (D) genera measured using Euclidean distance and “complete” method based on the relative abundances of differentially abundant genera between study groups.

The heatmap and hierarchical clustering of the differentially abundant bacterial genera revealed separated clusters of four sample types (Figure 5C). Interestingly, rhizosphere and bulk soil samples of both *Thymus* species were further clustered together. On the other hand, the heatmap and hierarchical clustering of the differentially abundant fungal genera revealed separated clusters according to the *Thymus* species (Figure 5D). The results indicate the variable characteristics of different kingdoms according to the soil sample type and plant species.

### 3.4. Predicted functional profile of bacterial microbiomes

To investigate the functional profile of the bacterial community in the soil samples, functional gene content was predicted and enumerated using Tax4Fun2. We used LefSe to determine differentially abundant KEGG functions (LDA>3) between rhizosphere and bulk soil samples for each *Thymus* species separately. The results revealed both overlapping and distinct functions separating rhizosphere and bulk soil samples in two *Thymus* species.

Quorum sensing, microbial metabolism in diverse environments, benzoate degradation, aminobenzoate degradation, degradation of aromatic compounds and sulfur metabolism were enriched in the bulk soils of both *Thymus* species. On the other hand, KEGG functions enriched in rhizosphere soils of the two *Thymus* species did not have any overlapping. We identified metabolic pathways, biosynthesis of secondary metabolites, biosynthesis of antibiotics, carbon metabolism, starch and sucrose metabolism, amino sugar and nucleotide sugar metabolism and Type I polyketide structures were enriched in the rhizosphere soil of *T. longicaulis* subsp. *chaubardii* (LS) (Figure 6A) while two-component system, biosynthesis of amino acids, porphyrin and chlorophyll metabolism and ribosome were enriched in the rhizosphere soil of endemic *T. pulvinatus* (Figure 6B).

**Figure 6.**
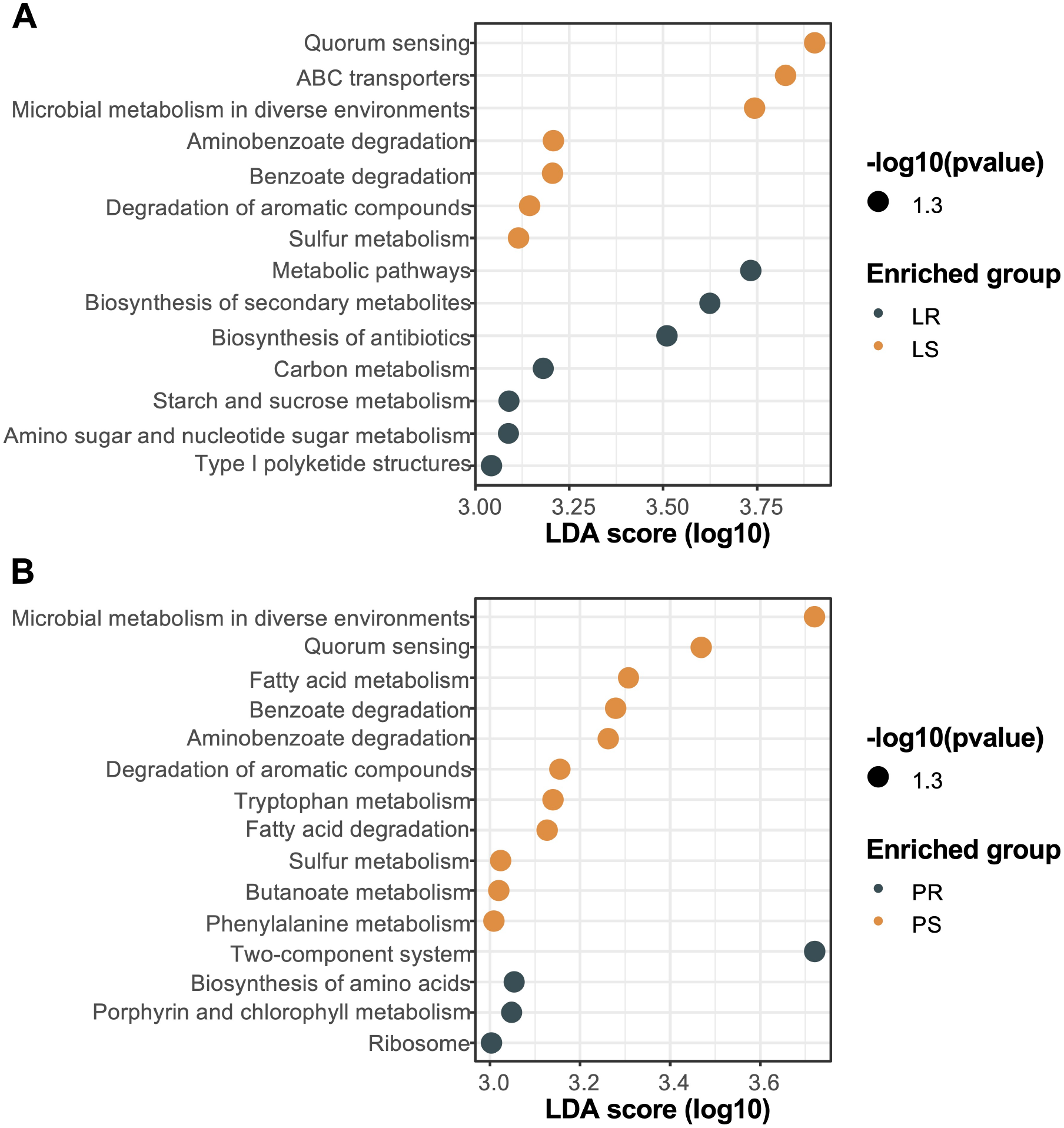
Tax4Fun2 predictions of the functional profile of *T. longicaulis* subsp. *chaubardii* (A) and *T. pulvinatus* (B) microbiome samples detected by LEfSe. LDA effect size (LEfSe) was calculated using LDA with p-value cutoff = 0.05 with LDA score > 3 of the genera. Dot sizes are indicative of -log10 (p value) while color is indicative of the study group with the enriched differentially abundant genera.

### 3.5. Integrative analysis of bacterial and fungal microbiomes

We used DIABLO to integrate 16S rRNA gene and ITS sequencing results and perform an exploratory examination of transkingdom interactions. The results revealed positive and negative correlations between bacterial and fungal ASVs in each *Thymus* species separately (Figure 7).

**Figure 7.**
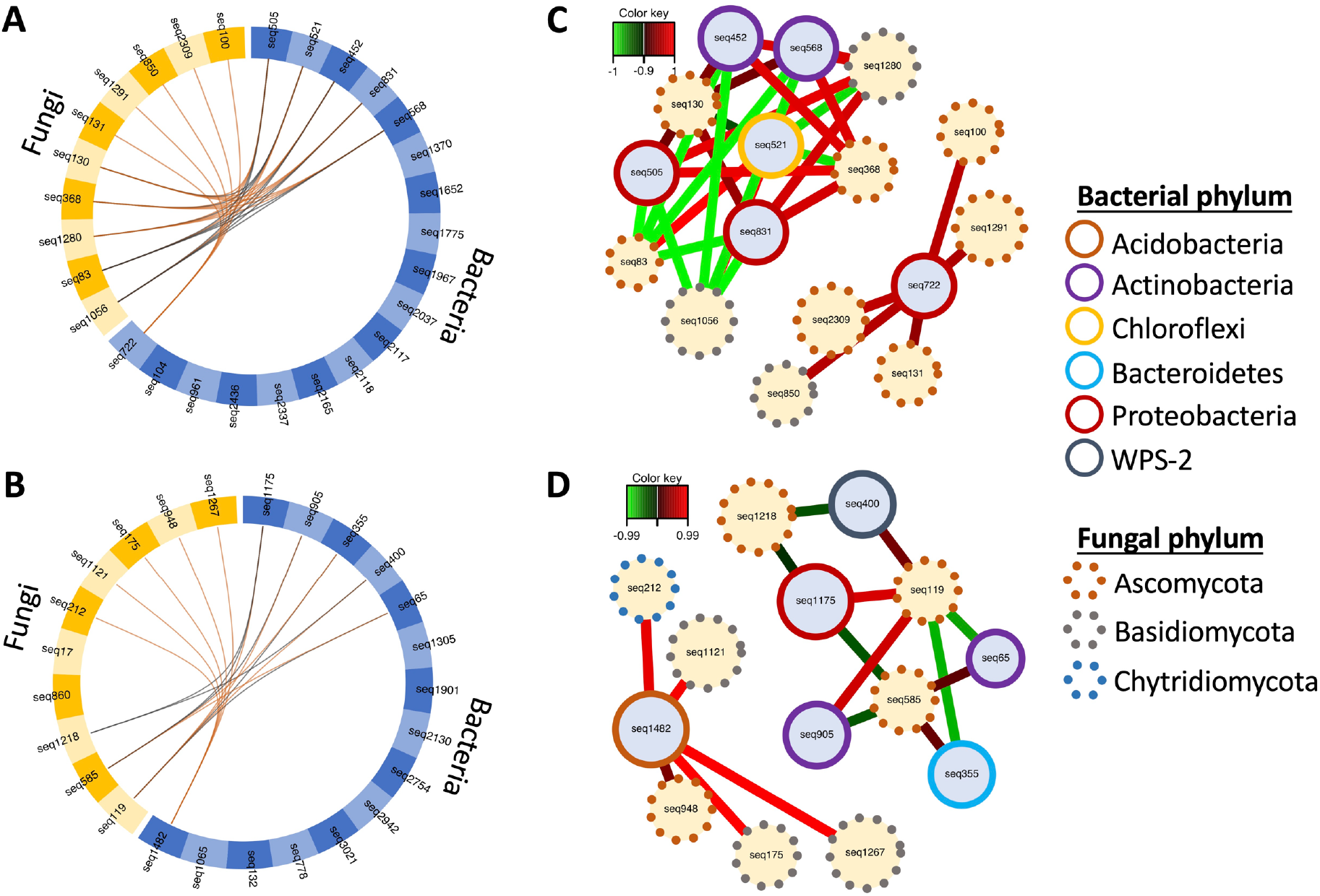
Circos plot representing the correlations between soil bacterial ASVs (blue side quadrant) and fungal ASVs (orange side quadrant) in *T. longicaulis* subsp. *chaubardii* (A) and *T. pulvinatus* (B). Positive and negative correlations (greater than 0.7) are illustrated with orange and black lines, respectively. Relevance network of bacterial ASVs and fungal ASVs in *T. longicaulis* subsp. *chaubardii* (C) and *T. pulvinatus* (D). Each node represents a selected ASV with the fill color indicating its type (light blue for bacterial ASVs and light yellow for fungal ASVs). Phylum level taxonomic assignments of bacterial and fungal ASVs are indicated with node line colors. The color of the edges represents positive or negative correlations as indicated in the color key.

In *T. longicaulis* subsp. *chaubardii*, we determined a strong positive correlation between bacterial ASV assigned to unclassified *Acetobacteraceae* and fungal ASVs assigned to *Didymella*, unclassified *Dermateaceae, Pseudolachnea, Cystobasidium* and *Alternaria* (Figure 7A). In *T. pulvinatus*, bacterial ASV assigned to *Bryobacter* was positively correlated with fungal ASVs assigned to *Sonoraphlyctis*, unclassified *Basidiomycota*, unclassified *Ascomycota, Rhizopogon* and *Tricholoma* (Figure 7B). Thus, bidirectional correlations between bacterial and fungal taxa were also species-specific. Altogether, our data show a bidirectional relationship between bacterial and fungal taxa that mutually influence each other in a species-specific way.

## 4. Discussion

In this study, we characterized the bacterial and fungal microbiomes of two *Thymus* species from Türkiye using 16S rRNA gene and ITS amplicon sequencing. We revealed that bacterial microbiome profiles differentiate not only plant species but also soil types (rhizosphere and bulk soils) while fungal microbiome profiles showed more distinct profiles between plant species. We identified discriminatory bacterial and fungal taxa between the plant species and soil types. Moreover, we showed the species-specific and overlapping functional profile changes between soil types. In addition, we applied an exploratory integrative approach and determined the species-specific nature of transkingdom interactions in two *Thymus* species.

Bacterial community profiles showed the *Proteobacteria, Actinobacteria, Acidobacteria* and *Chloroflexi* as the most abundant phyla in rhizosphere and bulk soil samples. In the previous study on the bacterial diversity in the rhizosphere of *T. zygis* growing in the Sierra Nevada National Park (Spain) reported *Proteobacteria, Actinobacteria, Acidobacteria*, and *Gemmatimonadetes* as the dominant phyla (26) which is consistent with our findings. In a previous study examining the fungal diversity of *Thymus* species, it was revealed that the cultured fungal isolates mainly consisted of the phylum *Ascomycota* (27). Similarly, our results revealed that *Ascomycota* was the dominant fungal phylum in our samples. Alpha diversity of bacterial and fungal communities did not show any significant differences between study groups while beta diversity analysis revealed significant differences between four sample types for both bacterial and fungal microbiome samples. However, it should be noted that the fungal microbiome samples mainly clustered based on plant species and clustering on the basis of soil type was less distinguishing. On the other hand, bacterial microbiome samples showed distinct clusters for each sample type indicating importance of both plant species and soil type on the bacterial community.

Differential abundance analysis revealed that unclassified *Acetobacteraceae, Sphingomonas, Blastocatella* and *Marmoricola* were the bacterial genera most strongly associated with the rhizosphere soil of endemic *T. pulvinatus. Acetobacteraceae* and *Sphingomonas* members are known to metabolize diverse nutrients and play an important role in nitrogen fixation which makes them abundant in plant roots (28, 29). *Sphingomonas* can also metabolize hydrocarbons with differently organized genes than genera of *Pseudomonas* and are found to promote plant growth under different abiotic stress conditions (30, 31). The higher abundance of *Blastocatella* has been previously associated to the high content of soil total organic carbon (32) and as a member of *Acidobacteria* to have a potential role in the turnover and stability of soil organic carbon (33). *Marmoricola* can produce leucine aminopeptidase and chitinase that provides stress resistance in plants (34). Also, *Canditatus* Udaeobacter is an important member of *T. pulvinatus* bulk soil samples, unlike other samples. *Canditatus* Udaeobacter, which is largely unexplored soil bacterium, has been reported to be responsible for the hydrogen cycle and show multidrug resistance (35). These findings mainly point to the association of *T. pulvinatus* with the bacterial genera that are contributors to carbon fixation, hydrogen cycling and nitrogen metabolism and provide stress resistance to the plants. Moreover, rhizosphere and soil samples belonging to *T. pulvinatus* had enriched Gram-negative bacterial genera while rhizosphere and soil samples belonging to *T. longicaulis* subsp. *chaubardii* were mostly associated with the increased abundance of Gram-positive bacterial genera, especially in rhizosphere soil samples. Gram-negative bacteria are known to use and be dependent on plant-derived carbon sources, while Gram-positive bacteria use carbon sources derived from soil organic matter (36). Based on this, Gram-negative bacteria enrichment in *T. pulvinatus* samples may indicate a higher plant-microbiome dependency. In contrast to bacteria, only a few fungal genera differed in relative abundance between study groups; *Saitozyma*, unclassified *Tylosporaceae, Geminibasidium, Solicoccozyma* and unclassified *Leotiomycetes* were enriched in bulk soil samples of *T. longicaulis* subsp. *chaubardii* (LS) while *Comoclathris* and unclassified *Chaetothyriales* were enriched in the rhizosphere soil of endemic *T. pulvinatus. Chaetothyriales* have been reported as plant symbionts and transmitted by seed (37). *Chaetothyriales* are capable of producing swainsonine which is an indolizidine alkaloid that causes severe toxicity in livestock feeding with swainsonine containing plants (38). Also, calystegines produced as a plant secondary metabolite are known to enhance the toxicity of plants with swainsonine which is only produced by endophytic fungi (39, 40). Because *T. pulvinatus* is a local endemic and critically endangered species, *Chaetothyriales* enrichment in its rhizosphere soil may be due to a protection mechanism. *Comoclathris* strains isolated from different plants and geographic areas are found to have a whitening effect by producing the same active metabolites. This metabolite, comoclathrin, is a tyrosinase inhibitor (41). Tyrosinase inhibitors are shown to be potential antibacterial agents (42) so the presence of *Comoclathris* may be involved in protection mechanisms or recruitment of bacteria to the roots of *T. pulvinatus*.

We compared the predicted functional profiles of rhizosphere and bulk soil types in two *Thymus* species separately to determine both differences and overlaps in enriched KEGG functions. Biosynthesis of secondary metabolites, biosynthesis of antibiotics, carbon metabolism, starch and sucrose metabolism, amino sugar and nucleotide sugar metabolism and Type I polyketide structures were enriched in the rhizosphere soil of *T. longicaulis* subsp. *chaubardii* while two-component system, biosynthesis of amino acids, porphyrin and chlorophyll metabolism and ribosome were enriched in the rhizosphere soil of endemic *T. pulvinatus*. The overlapping enriched functions of bulk soil samples have included quorum sensing (QS), microbial metabolism in diverse environments, benzoate degradation, aminobenzoate degradation, degradation of aromatic compounds and sulfur metabolism, whereas the rhizosphere soils of the 2 *Thymus* species did not have any overlapping enriched functions. In our results, contrary to the other studies, QS enrichment was in bulk soil instead of rhizosphere soil. Previous studies have indicated that the rhizosphere microbiome may vary depending on the different developmental stages of the plant (43–45) or season associated functional shifts (45, 46). The plant samples used in our study were collected near the end of the flowering period. Therefore, QS enrichment in bulk soil rather than rhizosphere soil may be due to the developmental stage of our plant samples. Moreover, two-component systems which are enriched in PR samples, are signal transduction systems that enable bacteria to sense, respond, and adapt to changes. Histidine kinases in quorum sensing mechanisms are part of two-component systems so although QS appears to be enriched in the soil, signal transduction systems are enriched in the rhizosphere of *T. pulvinatus*. Plants attract microbes through their tissues majorly by producing chemical signals and products such as amino acids. Presence of amino acids in the roots provides microbial community richness at the rhizosphere. It is known that microbial compounds can increase plant efflux of amino acids and contribute to microbial recruitment to the plant tissues (47). Enrichment of biosynthesis of amino acids in PR samples may be used to take up microbes from soil during different developmental stages and stress factors to provide plants survival. These functional profile differences indicate that the rhizosphere of endemic *Thymus* species mostly consists of specific bacterial groups that can function in plant protection and survival mechanisms while the rhizosphere of non-endemic species mostly consists of typical soil bacterial components.

We also evaluated bidirectional correlations between bacterial and fungal ASVs to identify a highly correlated transkingdom signature discriminating rhizosphere and bulk soil in each *Thymus* species separately. We identified a strong positive correlation between bacterial ASV assigned to *Acetobacteraceae* family of *Proteobacteria* and fungal ASVs assigned to *Ascomycota* and *Basidiomycota* in *T. longicaulis* subsp. *chaubardii* while another bacterial ASV assigned to *Bryobacter* family of *Proteobacteria* was positively correlated with fungal ASVs assigned to *Ascomycota, Basidiomycota*, and *Chytridiomycota* in *T. pulvinatus*. These findings indicate a bidirectional relationship between bacterial and fungal taxa that mutually influence each other in a species-specific manner. It should be noted that interaction related to *T. pulvinatus* samples was fewer and showed less complexity than *T. longicaulis* related interactions.

In conclusion, our study presents an overview of differences and similarities between the root associated microbiome profiles of two *Thymus* species, namely *T. longicaulis* subsp. *chaubardii* and *T. pulvinatus*. Supporting our hypotheses, our results provide a basis for future research on the plant-microbiome interactions in respect of plant endemism and contribute to the efforts of developing better conservation strategies for endemic plants.

## Acknowledgements

We thank Prof. Dr. Fatih Satıl (Balıkesir University) for guiding us to locate the plant species. This work was supported by the Scientific Research Projects Coordination Unit of Istanbul University (Project number: FYL-2022-39046).

## Author contributions

Conception and Design: GE, MA, FEÇY; Sample Collection and Processing: GE, MA, İSY, FEÇY; Data Analysis: GE, MA; Data Interpretation: GE, MA, FEÇY; Manuscript Writing – Original Draft: GE, MA, FEÇY; Review & Editing: GE, MA, FEÇY. All authors read and approved the final manuscript.

